# Evidence for predictive computations in a brain hierarchy during a visual search task

**DOI:** 10.64898/2026.04.09.717389

**Authors:** Dimitris A. Pinotsis, Andre M. Bastos, Earl K. Miller

**Affiliations:** Department of Psychology and Neuroscience, City St George’s—University of London, London EC1V 0HB, United Kingdom; Department of Psychology, Vanderbilt University, Nashville, TN, USA; The Picower Institute for Learning & Memory and Department of Brain and Cognitive Sciences, Massachusetts Institute of Technology, Cambridge, MA 02139, USA

**Keywords:** neural ensembles, attention, Predictive Coding, autoencoders, effective connectivity

## Abstract

Many lines of evidence suggest that the cortex functions fundamentally as a predictive system, compressing high-dimensional sensory inputs into low-dimensional representations that support internal models of the environment. These models generate top-down signals that shape and filter incoming sensory information, prioritizing unpredicted, and thus more informative, inputs for further processing. Multiple, mutually compatible computational frameworks have been proposed to explain these mechanisms, including predictive coding, predictive routing, and autoencoder-based approaches. However, a key challenge remains in empirically disentangling their relative contributions. Here, we directly compared three candidate algorithms, predictive coding, routing and autoencoders using LFP data from a brain hierarchy acquired during a visual search task. Notably, predictive coding differs from alternative models in its capacity to dynamically optimize signal passing across brain hierarchies in an input-specific manner, while the other algorithms do not consider constraints on feedback or feedforward inputs. It also goes beyond pairwise interactions, by predicting information flow in triplets of brain areas; here, V4, 7A and PFC. We considered the characteristic patterns of neural dynamics and within-area coupling that distinct algorithms produce. These were subsequently used to assess the relative evidence of each algorithm in the face of the LFP data. Our results support a hybrid account: hierarchical message passing consistent with predictive coding appears necessary to explain deep-layer activity, while predictive suppression mechanisms, aligned with predictive routing, account for superficial-layer dynamics without requiring explicit error computations.

## Introduction

Many lines of evidence suggest that the cortex is fundamentally a prediction system. Rather than deal with an overwhelming flood of sensory inputs, the architecture of the brain’s convergent anatomy reduces feedforward sensory-related neural activity to low-dimensional representations (Barrett and Miller, 2026; Pinotsis and Miller, 2022). This builds models of the world that can be used to automatically generate feedback signals that organize and filter and feedforward processing (Barrett and Miller, 2026; Friston, 2009; Pinotsis et al., 2017b, 2016a). The result is a filtering of predicted sensory inputs in favour of unpredicted (and therefore more informative) signals that are fed forward for full processing (Barron et al., 2020; Bastos et al., 2012; Bastos et al., 2020; Histed, 2025; Kanai et al., 2015). There are several, mutually compatible models and candidate mechanisms to explain this. Evidence for each exists. A challenge lies in assessing their relative evidence and contribution with empirical data.

Predictive coding models try to explain information flow in a hierarchy consisting of three or more brain areas and suggest that cortical circuits minimize prediction errors by comparing sensory inputs with top-down expectations (Friston, 2009; Pinotsis et al., 2017b). By modulating the gain of error computing neurons, a boosting of sensory channels that provide the most reliable information occurs (Parr and Friston, 2019; Pinotsis et al., 2014). This follows from theoretical arguments in Bayesian inference and has found some experimental support based on brain imaging data and computational modeling, see e.g. (Aitchison and Lengyel, 2017; Bastos et al., 2012; Feldman and Friston, 2010; Friston et al., 2015a; Huang and Rao, 2011; Millidge et al., 2021). It is also in accord with cortical function and anatomy (Barrett and Miller, 2026; Shipp et al., 2013). Predictive routing on the other hand, suggests that error signals are not necessary. Instead, prediction signals target neural representations of predicted sensory inputs and suppress them (Bastos et al., 2020a; Gabhart et al., 2023). The signals that are fed forward are those that were not suppressed. There are also active filtering mechanisms that can selectively amplify synaptic inputs of behaviourally relevant information via recurrent connections (Histed, 2025). Autoencoders on the other hand, suggest that state, not error, signals are fed forward—without feedback propagation (Hinton and Zemel, 1994). In short, different algorithms have been proposed to explain the same computations in cortical circuits.

Here, we adopt a new approach to directly compare the different models: First, we cast different brain algorithms, (predictive coding, autoencoders and predictive routing), into a common mathematical formalism –a General Linear Model (GLM). Next, we assessed the evidence of each candidate GLM using empirical data. This has the following advantages. First, it allows models to have the same number of parameters that can mitigate overfitting by a single model. Second, it highlights a key difference between Predictive Coding and other models: Predictive Coding dynamically optimizes neural wiring in an input—specific fashion. Third, it yields effective connectivity estimates that describe different paths that information might flow in a brain circuit (Pinotsis and Miller, 2022). This connectivity mirrors the different computations that might take place based on the brain algorithm implemented.

Using this approach, we compared the relative evidence of alternative brain algorithms. We used LFP data recorded from a brain hierarchy involving V4, 7A and PFC, during a visual search paradigm (Bastos et al., 2020a). The underlying hypothesis is that different brain algorithms would produce different neural dynamics and information flow in a brain hierarchy. This, in turn, will result in differences in neuronal coupling, that is effective connectivity. Differences in model evidence followed from these changes in effective connectivity and were quantified in terms of a Free Energy approximation. This is useful because the Free Energy penalizes for model complexity. Using the approach of (Pinotsis et al., 2017a; Pinotsis and Miller, 2023), we found the connectivity in brain circuits and built neural network models that implement different brain algorithms. By fitting these models to recorded LFP data, we then assessed the relative evidence of these algorithms. The evidence was in favour of a combination of predictive coding and routing models: State representations in deep layers seemed to require hierarchical message passing from areas above and below; at the same time, no explicit error computations seemed to take place in superficial layers and predictive suppression from deep layers offered a simpler explanation. This is an example of “computation through dynamics”, that suggests brain dynamics are the backbone of algorithm implementation (Driscoll et al., 2018).

## Methods

### Task and Experimental Setup

We analysed layer resolved LFP data recorded with laminar probes published in (Bastos et al., 2020a). Two rhesus monkeys (5—10 kg) were surgically implanted with head posts and recording chambers over V4, 7A and PFC. These were the only areas with layer resolved data needed for our analyses. Monkeys were positioned 50 cm from a 144-Hz LCD monitor (ASUS, Taiwan). Animals were trained using positive reinforcement to perform a visual search task (Figure 1A). Each trial began with fixation on a central point (2–3° radius) for 1 s, followed by presentation of one of three cue objects for 1 s. Monkeys were required to maintain fixation during a delay period of 0.5–1.2 s, after which a search array appeared containing the cued object together with one or two distractors presented at matched eccentricities (3–8°) but in different visual quadrants. Stimulus locations were randomly assigned on each trial. Correct saccades to the cued item were rewarded with juice. Behavioral performance was stable across animals (monkey S: 77% over 41 sessions; monkey L: 75% over 30 sessions). Training used a library of 22 images; for recordings we selected a subset of 12 images, with three objects used per session. In 65 of 71 sessions, the stimuli consisted of an orange, a green block, and a blue car. All procedures were approved by the MIT IACUC and followed the guidelines of the MIT Animal Care and Use Committee and the US National Institutes of Health.

**Figure 1.**
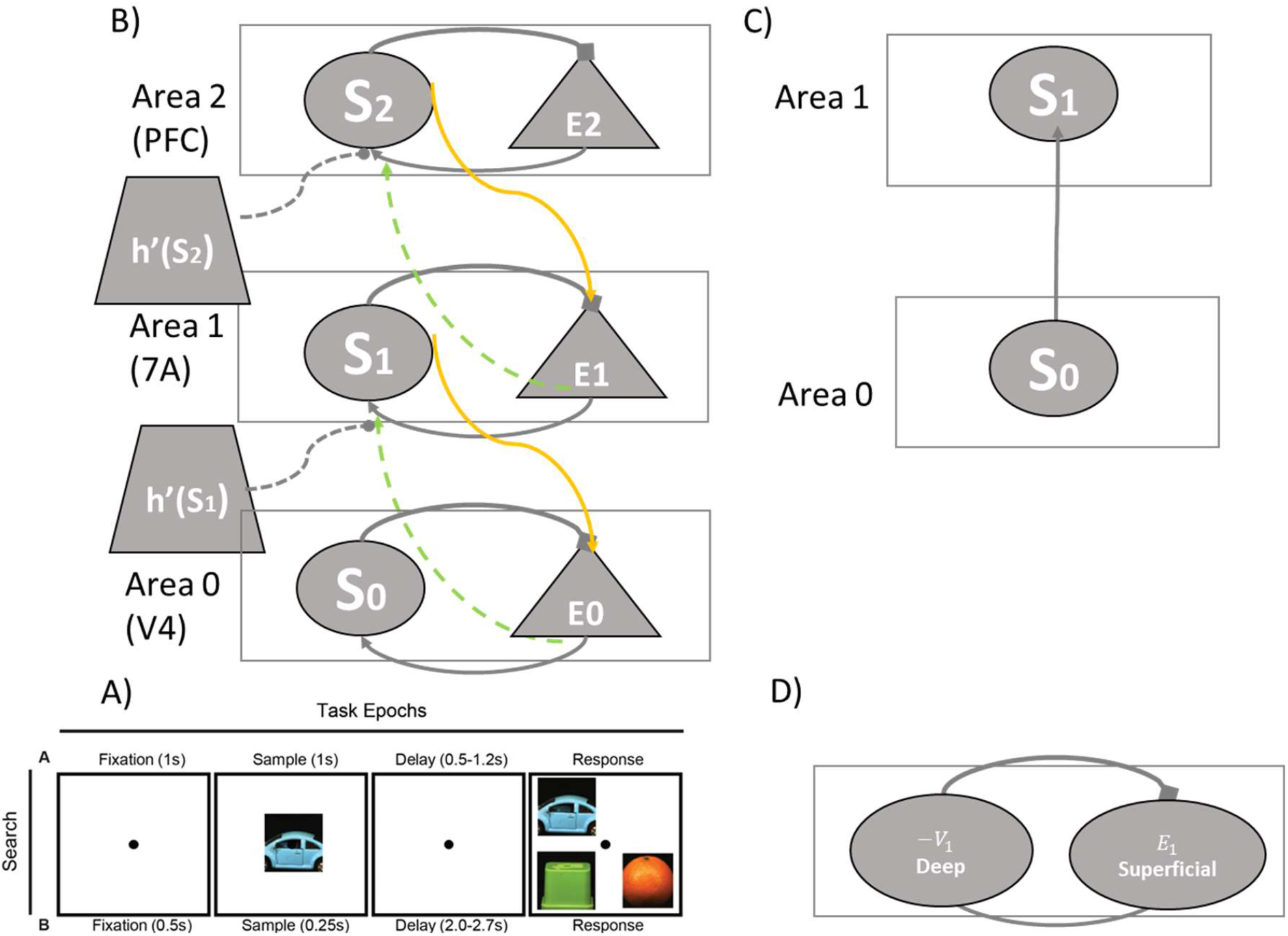
(A) Visual Search Task: After a one second fixation window, a sample stimulus (one of three pictures) is shown for one second. After a variable delay, the sample re-appears at one of 4 locations (always randomized). Monkeys saccade to the sampled stimulus. The sample identity was randomized (unpredictable blocks). Neural populations and message passing between brain areas according to Predictive Coding (B), autoencoders (C) and predictive routing (D). In Predictive Coding, state neurons (S_2_ disc) in the higher Area 2(PFC) send top-down expectations to error neurons (E_0_ triangle) in the lower Area 1(7A) (yellow arrow). Error neurons in Area 1 compute mismatches between predictions and sensory inputs and send error signals back up to Area 2 (dash green arrow). This input is scaled by the state output h(So)′ ∘ V^PC^ (see text). These propagate bottom-up, updating predictions iteratively, refining hierarchical perception and minimizing surprise across brain areas. Similarly, state neurons (S_1_ disc) in the higher Area 1 (7A) send top-down expectations to error neurons (E_0_ triangle) in the lower Area 0 (V4) (yellow arrow). Error neurons in Area 0 compute mismatches between predictions and sensory inputs and send error signals back up to Area 1 (dash green arrow). In autoencoders, state representations are passed up in the hierarchy from the lower population (S_0_ disc) to the higher state population (S_1_ disc)— without feedback propagation. Last, predictive routing includes the same state populations as Predictive Coding, oscillating at beta frequencies and occupying deep cortical layers as well as superficial populations oscillating at gamma frequencies. The difference from Predictive Coding is that the latter are not computing errors, but carry both stimulus and prediction information, see text for details.

### Learned connectivity in neural fields

In (Pinotsis et al., 2017a; Pinotsis and Miller, 2023), we used a neural field model (Amari, 1977; Coombes, 2007) to describe the transmembrane potential *V* in a group of neurons maintaining a memory, called neural ensemble. Neural field models describe the collective activity of large populations of neurons using a continuous, spatially distributed variable, denoted below by *z*. They model brain dynamics at a macroscopic scale by averaging neuronal activity over space and time, often using integro-differential equations. These models capture wave propagation, oscillations, and pattern formation in cortical activity. Applications include studying perception, decision-making, and brain disorders. Neural fields can be derived from a type of classical recurrent neural networks (RNNs) known as Wilson and Cowan equations (Grossberg, 2013; Wilson and Cowan, 1973) by replacing discrete neuron firing by neuronal currents produced by population activity that propagates on cortical patches. In (Pinotsis and Miller, 2022), we showed that starting from Wilson-Cowan Equations:

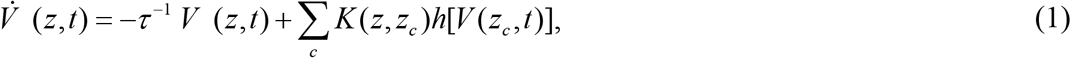

where *c* runs over all locations in the neural network except *z*, one can obtain replace the summation over *c* with an integral over space (a cortical patch, see below) and obtain a neural field

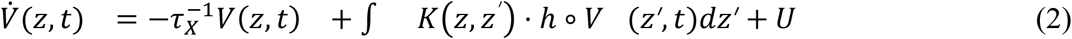

In the above equations, *z* is the location on the cortex from which the LFP electrode records data. In past work (Pinotsis and Miller, 2023), this was the location on a multielectrode array, while here this is the cortical layer from which each contact in our laminar electrode records. 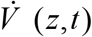 is the rate of change of the membrane potential at *z* and time *t*, 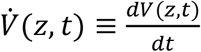, i.e. the temporal rate of change for an arbitrary function *h* is denoted by a dot above *h. K* is the connectivity matrix (i.e. a matrix whose entries weigh afferent input from location *z’* that arrives at *z*), *τ* is the time constant of postsynaptic filtering, 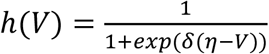 is a voltage to firing rate (transfer) function that describes population output, given its potential *V* and synaptic gain *δ*. Also, *η* denotes the postsynaptic potential at which half of the maximum firing rate is attained. The transfer function gives the instantaneous output of a neural ensemble with membrane potential *V*. Model parameters are included in Supplementary Table. To sum, Equation (2) describes the evolution of the potential *V* as a result of decay and recurring inputs from other parts of the neural ensemble. Further details and other applications of neural fields can be found in (Pinotsis et al., 2012) and references therein.

In (Pinotsis and Miller, 2023), we obtained the effective connectivity *K*(*z, z′*) after training model (2) with LFP data from a spatial memory task. This connectivity characterizes the weights by which a signal originating from location (or electrode located at) *z’*, is multiplied before reaching a neural population at location *z* –whose activity is measured with an electrode positioned at the same location. In other words, similarly to RNNs, we trained the neural field model using LFP data recorded using electrodes located at *z* and *z’* and obtained an effective connectivity that describes information flow in a cortical patch and the directed influence of one population on another. We called the neural field model with this connectivity, “*deep*” neural field to distinguish it from common neural field models where the connectivity is not learned but is chosen *ad hoc* and learning uses a “deep” (bottleneck) architecture.

The effective connectivity quantifies the causal influence of activity at *z’* on activity at *z*. This is similar to effective connectivity used in studies of large brain networks with e.g. fMRI (Friston, 2011; Van Den Heuvel and Pol, 2010) but at the scale of a few millimetres of cortical tissue. In (Pinotsis et al., 2017a), we used the above model to identify neural ensembles representing memories during a spatial memory task, while in (Pinotsis and Miller, 2022) we showed that these memory representations exist also at the level of the electric fields generated by these ensembles and are more stable than the corresponding representations at the neural population (spiking) level. Then, in (Pinotsis and Miller, 2023), we studied information exchanges between the electric field and neural levels. We found that electric field sculpts neural activity to tie together neurons in diverse areas that participate in memory ensembles. Here, the effective connectivity describes information flow between cortical layers sampled with laminar electrodes. This includes parameters in a deep neural field across depth as opposed to over the cortical surface considered in our earlier work. The model predicts information propagation during a sustained attention task (Srinivasan et al., 2023; Zhou et al., 2016). Our deep neural field model focuses on biophysics (i.e. time constant, couplings, activation function) underlying neural responses –but does not determine the algorithm the brain might be using to implement attention. As we will see below, various algorithms can be implemented using variants of the deep neural field model given by Equation (2). First, we consider Predictive Coding. After explaining what Predictive Coding is, we show how it can be implemented in our deep neural field discussed above.

### Deep neural fields for attention

To describe the algorithm the brain might be using to implement attention, we use Predictive Coding (Adams et al., 2016; Bastos et al., 2012; Friston et al., 2015a; Pinotsis et al., 2014; Rao and Ballard, 1999). This theory suggests that sensory information processing occurs while the brain samples its environment in a probabilistic fashion. This gives rise to stimulus representations in neural populations. It is an iterative process during which errors signals (discrepancies) between stimulus representations and sensory samples are computed at each time point and at each level of the visual hierarchy. Errors are used to update representations so that these get closer to the stimulus. As time goes by and the agent gathers more sensory samples, the corresponding errors reduce and representations become more faithful. Predictive Coding suggests that attention changes the update rate of representations so that they become more efficient, that is, they improve faster, in fewer sampling cycles.

The theory of Predictive Coding (PC) supposes that the brain is organized in a hierarchical fashion. Brain areas are thought to be functionally specialized and to form a core-periphery structure (Friston, 2008; Gu et al., 2019). PC suggests that sensory samples are inputs (from an area below in the hierarchy) and representations, that is state signals, follow from top-down prediction signals (i.e. input from the area above). To understand this, we consider the joint activity of brain areas 0 and 1, see Figure 1B. In this setting, area 1 (above area 0) sends top-down state signals to area 0, and area 0 sends bottom up state signals to area 1. The bottom up signals correspond to errors in area 0 generated by subtracting sensory inputs from predictions. The top down signals correspond to predictions from area 1. Similarly, area 2 (above area 1) sends top-down state signals to area 1, and area 1 sends bottom up state signals to area 2. In *Results* below, areas 0, 1 and 2 correspond to V4, 7A and PFC. Predictive Coding also suggests that error signals will be propagated in gamma frequencies up in the hierarchy and will be represented in superficial pyramidal cells from which feedforward connections originate predominantly (Bastos et al., 2012). At the same time, state representations will be sent down the hierarchy via feedback connections originating predominantly in deeper layers that also show beta frequency oscillations (Mendoza-Halliday et al., 2024).

To sum, Predictive Coding suggests that information flows in both ways in a brain hierarchy. It goes beyond pairwise interactions and predicts information flow in triplets of brain areas. Perceptual inference occurs while error and state signals propagate in opposite directions. State and error signals interact to minimize errors; these are computed in an online fashion. State populations denoted by *S* in Figure 1B, receive input from error populations denoted by *E*, in the same area and send state signals to the area below, while error populations send signals laterally and to the area above. Notably, state and error populations are thought to be oscillating in distinct frequencies, beta and gamma, and occupy different cortical depths (deep vs superficial). These spectral profiles have been confirmed using electrophysiology (Bastos et al., 2015; Miller et al., 2018; Vinck et al., 2022).

Having introduced Predictive Coding, we now present an extension of the deep neural field model (2) above that implements this algorithm. This extension will allow us to extend results from our earlier work on memory summarized above (Pinotsis and Miller, 2023, 2022) to the visual search paradigm considered below. For mathematical convenience consider now three random areas in a brain hierarchy: area *j+*1 and its two adjacent (higher and lower) areas, *j+*2 and *j*. The bidirectional information flow of state and error signals between areas *j, j+1* and *j+2* can be described by the following Equations

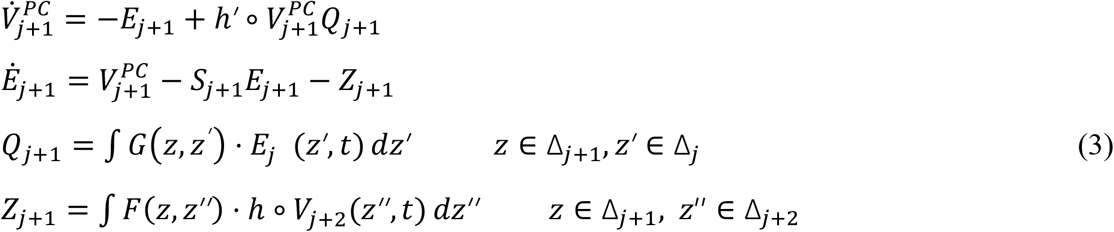

These equations describe average firing rate for the prediction 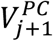 and the error populations *E*_*j*+1_. They are similar to the well-known Jansen and Rit (Jansen and Rit, 1995) and Dynamic Causal Modeling approaches (Marreiros et al., 2008; Pinotsis et al., 2017b) and are shown in Figure 1B. The difference between Equations (3) and those earlier models is that these equations included connections that are learned from the data similarly to neural network training. These describe the heterogeneous connections between the error and stare populations given by the matrices *G*(*z, z*^*’*^) and *F*(*z, z’’*). These learned weights are then used as priors for fitting LFP data, see *Results* and (Pinotsis and Miller, 2023) where a similar approach was used to test for ephaptic effects. The derivation of Equations (3) is included in the Supplementary Material. Below we discuss the meaning and implications of these Equations.

In Equations (3), *Δ*_*j*_ is the cortical patch containing the neural populations from where we sample in area *j*, and similarly for areas *j+1* and *j+2*. Locations within patches in areas *j, j+1* and *j+2* are parameterized by variables *z’, z* and *z* ***‘***. The remaining notations and functions appearing in Equations (3) are explained below, see also Figure 1B. Note that Equations (3) are formally similar to Equations (2) above, that is, they are deep neural fields. This will be explained below. We call Equations (3) *deep neural field equations for attention*. The first line in Equation (3) is formally similar to Equation (2). In Equation (3), the decay term on the right hand side (RHS) includes lateral input in the form of error activity (1^st^ term on RHS; shown by a thick line terminating with a arrow from E_1_ to S_1_ in Figure 1B), while in Equation (2) the decay term included state, not error, activity. Further, in Equation (3) there is an additional error input *Q*_*j*+1_ (see the third line of Equation (3)), that arrives to the state population from the area *j* below (weighted by feedforward connectivity *G*(*z, z*′), see also the dashed green line from E_0_ to S_1_ in Figure 1B). This input is scaled by the state output *h*(*So*)*′* ∘ *V*^PC^. This has a multiplicative effect over incoming error signals from the area below. In the case of a sigmoid transfer function *h* considered here (see Equation (1)), the derivative *h′* corresponds to activity variance across the population (Marreiros et al., 2008), denoted by a trapezoid in Figure 1B. This is the first instance of “precision-weighting” that is a central tenet of Predictive Coding. Precision is defined as the inverse variance of activity across the population. In other words, error signals are weighted by precision (Pinotsis et al., 2016b, 2014). If precision improves, errors are more efficient in constructing representations. Precision weighting is thought to result from neuromodulation. According to some authors, this is the main mechanism for implementing attention, see e.g. (Feldman and Friston, 2010; Friston et al., 2012). We now consider another instance of precision weighting in Predictive Coding by focusing on the second line in Equation (3):

This describes the temporal evolution of error populations. According to this equation, errors are computed based on the difference between local state responses 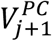 (see line from S_1_ to E_1_ terminating with a square in Figure 1B) and top down state input (predictions) *Z*_*j*+2_, weighted by feedback connectivity *F*(*z, z*′) (see the last line in Equation (3) and yellow line from S_1_ to E_0_ in Figure 1B). The second line in Equation (3) also contains a decay term, with rate equal to the precision 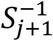 of the error population. This is precision weighting: the error populations reach a steady state at a rate determined by the population precision. The larger the precision (or neural gain), the sooner the error populations reach the steady state. Error populations that carry reliable information will also evolve faster than populations with less trustworthy information. Behaviourally, this is similar to longer decision times in noisier environments during sensory perception (Kriener et al., 2017; Vaden Jr et al., 2022).

This is the second instance of “precision-weighting” suggested by Predictive Coding. The first was the weighting of error input from the area below discussed above. Overall, precision weighting implements a mechanism by which the brain selects error representations in the light of incoming sensory evidence: the more reliable this is, the faster error populations update state representations. This is reminiscent of Kalman filtering, where precision corresponds to Kalman gain (Adams et al., 2016). Precision weighting allows the brain to weight upstream errors in an online, moment by moment fashion, so that more noteworthy information is used to sharpen states, and reduces errors faster in the next iteration.

To sum, Equations (3) are dynamic equations for state and error population activity. This is the activity one measures in populations in areas *j, j+1* and *j+2* –using e.g. local field potential (LFP) electrodes—assuming that these populations compute state and error signals. Following a large body of work on PC (Barron et al., 2020; Bastos et al., 2012; Gabhart et al., 2023; Kanai et al., 2015) we here associate state or predictions signals wit LFPs from deep cortical layers and error signals with superficial layers and consider recordings from areas V4, 7A and PFC cortex and ask whether this associations has sufficient evidence in light of the data, see *Results*.

### Alternatives to Predictive Coding

Above we considered Equations (3) that predict neural activity during an attention task. Implementing attention was achieved via Predictive Coding. Equations (3) include distinct Equations for state and error populations (first and second line). These were derived starting from the premise that information about states is propagated with minimal loss between areas, see Supplementary Material for more details. A well known alternative to Predictive Coding is offered by the theory of Autoencoders (Hinton and Zemel, 1994; Linsker, 1988). This suggests that stimulus information is preserved locally at each area of a brain hierarchy. The neural population carrying this representation is driven by upstream sensory input from population S_0_ to S_1_, see Figure 1C. We define a probability density that describes stimulus processing while the brain samples the visual field

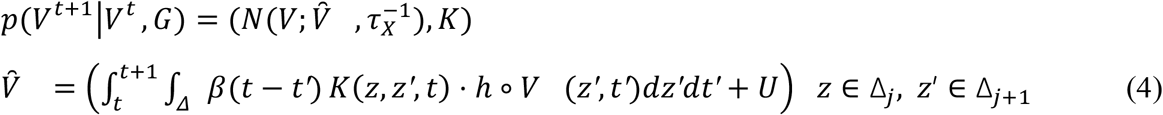

This is an example of a generative density that describes the probability with which neural activity *V* is generated at each time point given a recurrent (intrinsic) connectivity *K*(*z, z′, t*) and a postsynaptic time constant *τ*_*X*_. Generative densities appear in the theory of representational learning discussed in the next section. In brief, they provide a mapping from parameters (like time constants and connectivity) to observed data. The function 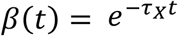 describes the time course of post synaptic potentials (Rall, 1967). Here and below, *N*(*x*; *m, S*) is a normal distribution over a variable *x* with sufficient statistics *m* and *S*; we assumed the generative density follows a normal distribution; something known as a Laplace approximation (MacKay, 2003). This is a common assumption in population coding following from the central limit theorem (Knill and Pouget, 2004) The second line in Equation (4) incorporates spatial and temporal summation. It suggests that the membrane potential is the result of synaptic inputs from across the population and decaying activity from an earlier time point *t* until the current time, *t*+1, and external input, *U*. This is the reverberating activity (output) of the neural ensemble during the epoch defined by beginning and end of the interval [*t,t*+1] mediated by recurrent connectivity *K*(*z, z′, t*). Note that, for mathematical simplicity, we have neglected local conduction delays.

The first line in Equation (4) gives the probability that the population fires at *t*, given a history of synaptic inputs across the population. In this setting, the goal of this firing is the accurate stimulus (state) reconstruction. The corresponding loss is simply taken as the sum of (log) probabilities across all time points *t*=1,…*T* 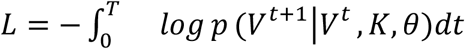. In other words, maintaining stimulus information, for each time point during the visual search task implies that the probability *p*(*V*^*t*+1^|*V*^*t*^, *θ*) is maximum, that is, state information is reconstructed accurately at each time point. Then, using gradient descent 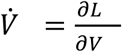, we can find the equation that describes the temporal evolution of neuronal activity: this is given by a deep neural field,

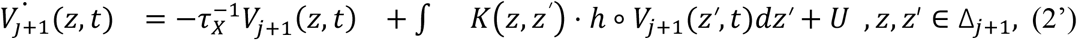

The above equation is similar to Equation (2) that describes a deep neural field.Thus we call this Equation (2’). As before, the locations on the cortical patch in area *j* are parameterized by *z* and *z’*. To sum, starting from an optimization goal (maintain stimulus information) we derived the equation for neural activity in autoencoders. The corresponding proof for Predictive Coding is similar and is included in the Supplementary Material.

Besides autoencoders, another alternative to Predictive Coding, is predictive routing (Bastos et al., 2020a). Similarly to Predictive Coding, predictive routing assumes there are prediction (state) neurons (oscillating in beta frequencies). The difference is there are no error neurons, see Figure 1D. This is motivated by recent studies using global and local oddballs (Gabhart et al., 2023) and oddball responses in the absence of top-down predictions(Xiong et al., 2024). These conclude that pyramidal neurons do not necessarily compute errors. There is feedback from higher to sensory brain areas – but this often does not affect spiking. Mathematically, we express the activity of predictive routing neurons situated in superficial cortical layers by an Equation similar to Equation (2’), with an additional inhibitory drive by deep layer activity, *V*_*j*+1_ representing state activity,

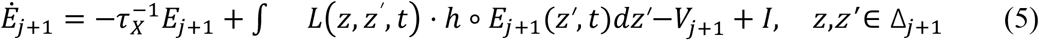

This mirrors the circuitry presented in Figure 1D. Here, *L* is the recurrent connectivity of the superficial pyramidal cell population *E*_*j*+1_ (Figure 1D) and *I* includes extrinsic input to the pyramidal cell population from other brain areas, while *V*_*j*+1_ is the activity of the deep pyramidal cell population (Figure 1D). This is the predictive routing (PR) model that we use in *Results*.

### Learned connectivity reflects the brain’s algorithm

How can we identify which algorithm the brain might be implementing? Here we suggest that an answer is obtained by first computing the effective connectivity of neural circuits that are implementing various candidate algorithms. Having obtained the connectivity, we can then obtain predictions of brain activity using a neural network that contains this connectivity. Finally, we score the relative evidence of these predictions by comparing them against recorded LFPs.

The three algorithms considered above suggest different ways that information is passed around within cortical circuits. For example, Predictive Coding suggests errors are computed and exchanged, while the other algorithms do not. Also, autoencoders suggest that information flows one way, i.e. bottom up, while the other algorithms consider a bidirectional flow. Such differences are built in the model Equations (2’), (3) and (5). In *Results*, we train the three models using LFP data from an attention task (Bastos et al., 2020a) and the approach of (Pinotsis et al., 2017a; Pinotsis and Miller, 2022). We thus obtain the learned connectivity, i.e. after the animal has undergone behavioral training. This is the connectivity in neural populations that implement Predictive Coding, autoencoders and predictive routing. But how can we be sure that this connectivity can be learned? To see how, we need to discuss the theory of representational learning (Dayan and Abbott, 2005).

Recall that earlier, we defined a generative density for autoencoders *p*(*V*^*t*+1^|*V*^*t*^, *K, θ*) –see Equation (4). This yielded neural activity conditioned upon some model parameters (e.g. connectivity *K* and other model parameters *θ*). Then starting from the generative density *p*(*V*^*t*+1^|*V*^*t*^, *K, θ*), we obtained equations for neural activity (Equations (2’)). The same process can be followed for other brain algorithms. For Predictive Coding, we define the generative density, as follows

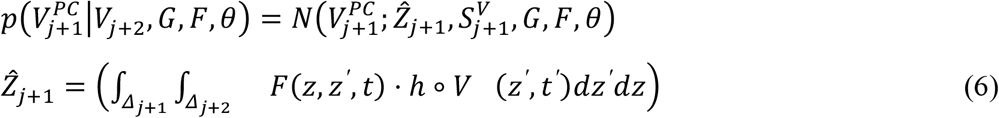

Here 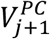 is the firing rate of the state equation representing stimulus information in area *j+1*. The Greek letter *θ* is used to lump together model parameters. Starting from the probability density of Equation (6), one can obtain the Predictive Coding Equations for neural activity (3), see Supplementary Material. This is mathematically similar to the proof in the previous section where we derived the Equations for autoencoders (2’). In both cases, optimizing the generative density yields the neural equations. In autoencoders, this optimization suggested the conservation of stimulus information as it propagates upstream. In Predictive Coding, the optimization of the generative density (6) suggests that the error, i.e. the divergence between top down predictions and bottom up inputs is minimum. Further, the density (6) connects the feedforward and feedback connectivity *G, F* to neural activity *V* = {*V*_*j*+1_, *V*_*j*+2_}.

Having obtained the generative density, the theory of representational learning suggests that one can also define the recognition density. This is complementary to the generative density considered above: it yields the model parameters, (e.g. connectivity *G, F*) conditioned upon neural activity (e.g. LFP recordings). The correspondence between generative and recognition densities follows from Bayes rule; the recognition density is given in terms of the generative density (Dayan and Abbott, 2005):

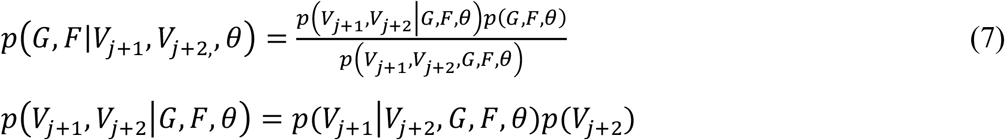

Both the generative, i.e. Equation (6), and recognition, i.e. Equation (7) densities are used in representational learning. Crucially, the theory of representational learning suggests that after learning is completed, neural dynamics that are given by Equations (2’) and (3), will have reached a steady state. This will be used below to simplify Equations (2’), (3) and (5). At steady state, the mean and variance of the recognition density *p*(*G, F*|*V*_*j*+1_, *V*_*j*+2_, *θ*) can be obtained from their stationary values 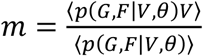 and 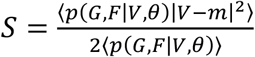. In these formulae, brackets denote averages over all sample values, 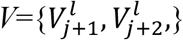, and *p*(*G, F*|*V, θ*), is a shorthand for the recognition density, see also (Dayan and Abbott, 2005). Thus, posterior or recognition densities for the connectivity follow naturally from learning. This mirrors our experimental intuition that connectivity is shaped by learning using plasticity mechanisms (Caporale and Dan, 2008). To sum, representational learning suggests that the connectivity *G* and *F* (as well as *K* and *L*) can be obtained from LFPs after training. Examples of *G, F, K* and *L* will be presented in *Results*.

### Learned connectivity in autoencoders and predictive routing

Having justified *why* connectivity can be learned, below we explain *how* we implemented learning and found the connectivity. We used the approach of our earlier work (Pinotsis and Miller, 2023, 2022). This is summarized below. The connectivity *K*(*z, z*^*’*^, *t*) of a neural field model given by Equation (2) was learned by reformulating a deep neural field as a Gaussian Linear Model (GLM) and using a Restricted Maximum-Likelihood (ReML) algorithm (Harville, 1977). The advantage of this neural field model is that it includes learned effective connectivity, that corresponds to the connectivity in a neural circuit of the animal that has learned to perform the task. We call it a *deep* neural field model to distinguish it from other neural mass and field models where connectivity is prescribed ad hoc.

Below we show that we can find the connectivity in three deep neural fields. These implement the brain algorithms considered above: Predictive Coding, autoencoders and predictive routing. Despite differences in the way information is passed around in a cortical circuit, all three algorithms can be written in the generic form of a deep neural field, i.e. Equation (2). We show this below. The difference will be in the recurrent connectivity *K*(*z, z′, t*). We first consider the connectivity *K*(*z, z′, t*) in an autoencoder model. This is the simplest among the three models. We saw above that during learning the model converges and the mean and variance of neural activity have reached their stationary values (steady state; Dayan and Abbott, 2005). Stationarity implies that 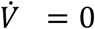 and Equation (2’) for the neural depolarization yields

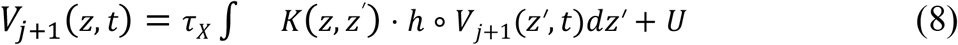

Since neural activity has reached a stable (stationary) value we neglect the temporal dependence in the connectivity and simply write *K*(*z, z′, t*) = *K*(*z, z′*). Then, training the above neural field model with LFP data *Y*_*j+1*_ from area *j+1*, we obtain a prediction of neural activity in the Autoencoder (see Supplementary Material for details). This activity is given by the following GLM:

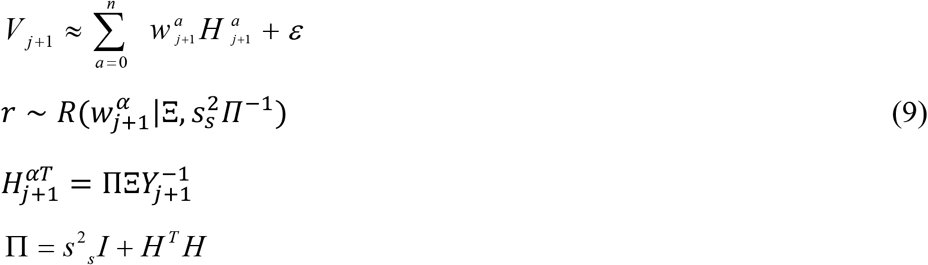

where we assumed that the connectivity components 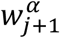 can be sampled from a normal prior distribution 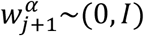 and *ε* : (0,*σ* ^2^ I). This is the neural activity in the autoencoder model (8)—reformulated as a GLM, i.e. a product of connectivity components *w*^*a*^ times principal axes 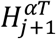. The principal axes 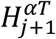, defined in the third line of Equation (9), describe the temporal evolution of neural activity. They are given by the spatial derivatives of the depolarization 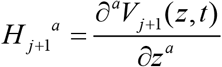,They have dimension *N*_*T*_ *x T*, trials by samples, and their values are equal to the instantaneous scale factor with which the corresponding connectivity 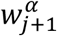 must be multiplied to reconstruct the observed LFP. They predict the oscillatory profile of fluctuations around baseline. These fluctuations can be thought of as transient non Turing patterns (patterns that decay back to baseline) whose Lyapunov exponents determine the frequencies observed in sampled LFP activity. The second line in Equation (9) suggests that the connectivity 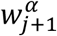 is determined by the posterior mode, *Ξ*,of the approximate posterior *r* to the true posterior *p* ~N(*w* |*Y*). The connectivity components 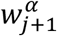 have dimensions equal to the number of electrodes by the number of trials and are an alternative characterization of the intrinsic connectivity *K* : knowing 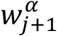, uniquely determines *K*. They describe how signal exchanged between different neural populations (in the vicinity of each electrode) is weighted as information is transmitted between different cortical layers.

To sum, following (Pinotsis and Miller, 2022) we find the connectivity *Ξ* as the mean of an approximate posterior distribution *r* with variance 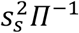. Then, the principal axes 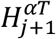 are given by the third line in Equation (9) and neural activity in a neural circuit (network) that includes this connectivity, is given by the first line in Equation (9).

Above, we described how (effective) connectivity can be learned in a neural circuit that implements autoencoders. We now consider the predictive routing equations i.e. Equation (5). We derive the analogue of Equation (9) starting from the deep neural field (5). Learning the connectivity in a neural circuit that implements predictive routing is very similar to the autoencoder considered in the previous paragraph. This is because Equation (5) is also a deep neural field model, similar to the deep neural field for the autoencoder given by Equation (2’). Thus, we can construct the analogue of Equations (9) for predictive routing just by mimicking the autoencoder derivation. Again, starting from steady state assumptions (as in the autoencoder model above), *Ė* = 0, and we obtain

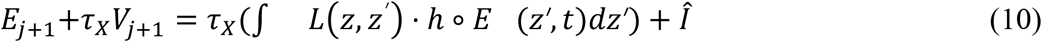

Recall that *V*_*j*+1_ is the activity of the deep pyramidal cell population. Then Equation (10) is the same as (8) where *V*_*j+1*_ is replaced by *E*_*j*+1_+*τ*_X_*V*_*j*+1_ and the exogenous input *U* is replaced by *Î* = *τ*_*X*_*I*. Then, the activity of the predictive routing model is given by an equation similar to the first line of Equation (9), after subtracting *τ*_*X*_*V*_*j*+1_

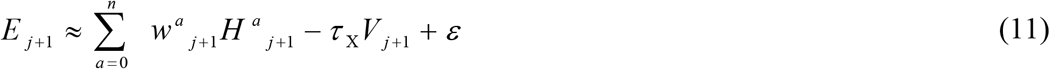

The quantities appearing in Equation (11) are given by lines 2—4 of Equation (9). The difference between the predictive routing and autoencoder models is that the former was trained with data from superficial LFP contacts, where the error population *E*_*j*+1_can be found. The autoencoder on the other hand, was trained with data from deep LFP contacts, where the state population implementing the autoencoder resides. In *Results*, we compute predictions for the activity of autoencoder and predictive routing models using Equations (9) and (11).

### Learned connectivity in Predictive Coding

Turning to Predictive Coding, we first consider the third and fourth lines of Equation (3) that predict the feedforward and feedback inputs *Q*_*j*+1_ and *Z*_*j*+1_: their form is identical to deep neural fields, given by Equation (2). Thus, in the same way we derived Equation (9), we obtain the following expressions for *Q*_*j*+1_ and *Z*_*j*+1_

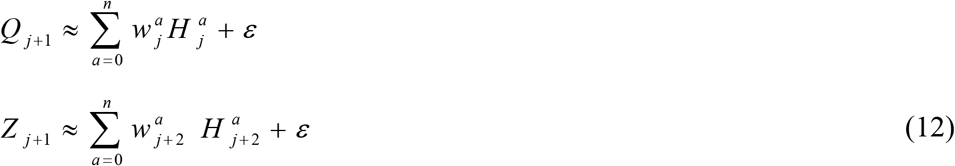

The only difference of the above equations from the autoencoder model above is that to find the feedforward input *Q*_*j*+1_, we trained the model with LFP data from the area *j* below (i.e. bottom area). This is indicated by the subscript in 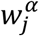. Similarly, to find the feedback input *Z*_*j*+1_, we trained the model with LFP data from the area *j+2* above (i.e. top area), see the subscript *j+2* in 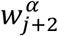. Briefly, the difference from previous models is that Predictive Coding considers the online, continuous message passing between (three) adjacent areas in a brain hierarchy as opposed to autoencoders and predictive routing that consider local dynamics. LFPs from all three areas were used for training as opposed to data from one area in the other two algorithms. The quantities *Q*_*j*+1_ and *Z*_*j*+1_ found above will be used below. Once *Q*_*j*+1_ and *Z*_*j*+1_ *are* known, Equations (3) yield predictions of state activity 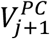. To see that, note that following stationarity (as above), 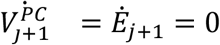 and Equations (3) yield

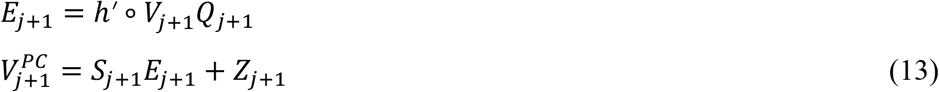

Eliminating *E*_*j*+1_ and combining the two Equations above, we obtain an expression for the state activity 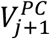, in terms of *Q*_*j*+1_ and *Z*_*j*+1_:

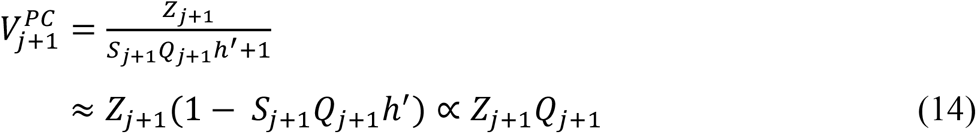

where *Q*_*j*+1_ and *Z*_*j*+1_ are given by Equations (12). To obtain the second line in Equation (14), we used the identity 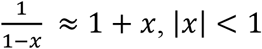 with *x* = −*S*_*j*+1_ *Q*_*j*+1_ *h*^′^ and absorbed constant terms in the connectivity. Equation (1) 4 motivates the following expression for the state population

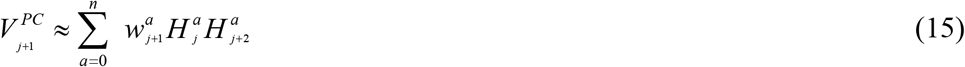

Equation (15) above suggests that activity in the Predictive Coding model 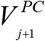, is given by a product of connectivity *w*^*a*^ times principal axes *H* (Pinotsis and Miller, 2022). In the derivation of Equation (15), our goal was to find an equation similar to Equation (9) above. We knew that the main difference of Predictive Coding from other brain algorithms is that connectivity is optimized online and in an input specific fashion, i.e. it is not defined by the prior distribution 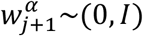, as in autoencoders above. Instead, there is an additional constraint that the principal axes 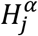 have an input specific (empirical) prior given by 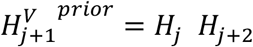. This is the product of the corresponding principal axes of the two areas that are adjacent to the area where the state population is located. In other words, the main difference of Predictive Coding from the autoencoder and predictive routing models above is that the latter models employ a bottleneck architecture (with independent Gaussian priors), while the former employs empirical priors, see. e.g (Hohwy et al., 2008). In the latter models, the recognition model is implemented in the connections from latent states to model predictions. The corresponding weights are found by maximizing the mutual information between latent states and model input. On the other hand, Predictive Coding utilizes some dynamic constraints that help solve the inverse problem of reconstructing the recognition density (Friston, 2009). These are given in the form of empirical priors (Friston et al., 2015b; Pinotsis et al., 2016c). To sum, the optimization of the cost function is carried out in an input specific fashion. This input consists of the product of inputs *H*_*j*_ from the area below and *H*_*j*+2_ from the area above. By repeating the derivation for the error population (instead of the state population considered above), we obtain

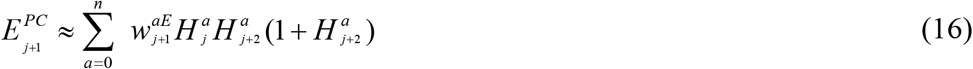

This follows from Equations (3) and stationarity. Equations (3) yield *E*_*j*+1_ ∝ *Z*_*j*+1_ *Q*_*j*+1_ (1+*Q*_*j*+1_) and the empirical prior is given by 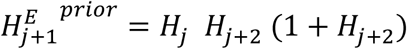. In *Results*, we compute predictions for the state and error activity of the Predictive Coding model using Equations (15) and (16).

## Results

### Effective connectivity across cortical layers in a visual search task

We analysed data recorded using laminar electrodes in three brain areas: area V4, 7A and PFC during a visual search task (*Methods*; Bastos et al., 2020). Note that although (Bastos et al., 2020) contains data from additional areas, this is the only hierarchical triplet of areas, for which LFP data were layer resolved. Having layer resolved data is necessary for all our analyses below, thus, these are the only areas we considered from that dataset. Also, having data from a brain hierarchy consisting of three areas is necessary for testing the integration of bottom up and top down signals suggested by Predictive Coding (see *Methods* and below).

Animals had to saccade to the sampled stimulus (Figure 1A). We assumed that these brain areas are connected to each other with feedforward and feedback connections: area 7A receives feedforward input from V4 and feedback input from PFC. Thus, 7A was assumed to be the middle area in a brain hierarchy with V4 being the area below and PFC the area above. We considered three models of neural activity: a Predictive Coding (PC) model, a predictive routing (PR) model and an autoencoder (AE) model (see *Methods* for details). The main difference between the PC model and the other two is that in the PC model the brain computes the error between its predictions and sensory input, while in the latter models do not. Also, the PC model describes how activity varies in a brain hierarchy of circuits in adjacent brain areas, while the other models do not offer such descriptions. We review these three models below.

In the PC model, an agent gathers sensory samples online to construct stimulus representations (Friston, 2009, 2008; Kanai et al., 2015). This is an iterative process in which the error (divergence) between the current representations and incoming sensory samples are computed at each time point. The error is then used to update the representations, gradually aligning them more closely with the actual stimulus. Over time, as the agent collects additional sensory samples, the error diminishes, and the representations become increasingly accurate. This imposes real time constraints (known as empirical priors (Friston, 2005)) on neural activity from other areas in the brain hierarchy –we will come back to this below.

PC further suggests that attention effects involved in the visual search task considered here will result in modulatory inputs that change the update process: The updates become more efficient and representations improve more quickly, that is, in fewer iterations. A summary of these ideas is depicted in Figure 1B using a trapezoid that weights incoming error signals from the area below. The theory of Predictive Coding proposes that signals are exchanged between neighbouring areas in a brain hierarchy. In Figure 1B, two such neighbouring areas are shown, Area 0 and Area 1. These could be any of the two pairs formed by the areas above, that is, V4 and 7A or 7A and PFC. They exchange signals via reciprocal connections, encoding state (i.e. prediction) and error signals, that interact to minimize errors. In sum, Predictive Coding suggests that sensory samples serve as inputs from lower areas, while state representations are shaped by top-down predictions from higher areas.

To understand this, consider the interactions among three brain areas, V4, 7A and PFC shown in Figure 1B. Each area comprises two populations that perform Predictive Coding: a pyramidal cell population representing errors (E_0_, E_1,_ and E_2_ depicted with triangles) and another population representing predictions or states (S_0_,S_1_ and S_2_ depicted with discs). Area 1 (7A) sends top-down predictive signals to area 0 (V4; yellow arrow), while area 0 provides bottom-up sensory inputs to area 1 (dash green arrow). These bottom-up inputs are compared against predictive signals, generating error signals at area 0. Similarly, area 2 (PFC) sends top-down predictive signals to area 1 (yellow arrow), while area 1 provides bottom-up sensory inputs to area 1 (dash green arrow).Thus, information flows bidirectionally within the brain hierarchy with state signals propagating top-down and error signals propagating bottom-up. This suggests that to test Predictive Coding one needs data from a hierarchy of three or more areas of the sort considered here. In the middle area 1 (7A) bottom up and top down signals from the other two areas are integrated and are passed again in higher and lower areas of the hierarchy.

Autoencoders (AE) are a simpler, alternative model to Predictive Coding (Figure 1C; Baldi, 2012; Hinton and Zemel, 1994). Here, information flows in a unidirectional fashion, like in a deep convolutional neural network. In this setting, each brain area comprises a single pyramidal population representing states (S_0_ and S_1_). The goal is to minimize the difference between the sensory input and the corresponding state representation. In autoencoder models, state signals are propagated upstream cortical hierarchies in a feedforward fashion. Note that the difference between Predictive Coding and autoencoders is twofold: first, Predictive Coding requires feedback signals –that are known to be important for conscious awareness. These are absent in autoencoders. Instead state signals are propagated in the reverse direction, that is, feedforward in autoencoders. This is the second difference.

In predictive routing (PR), predictions are formed in the deep layers and serve to inhibit gamma activity in superficial layers, see Figure 1D (Bastos et al., 2020; Gabhart et al., 2023). When a prediction—conveyed via deep-layer beta rhythms—suppresses the incoming sensory input, the result is reduced superficial-layer gamma activity. This looks like an absence of “prediction error” signal in the superficial layers. Note that PR does not need a dedicated error-computation process in layer 2/3 neurons. Rather, it describes some feedforward activity in the absence of predictive modulation. In other words, layer 2/3 activity is not a specialized error signal, but rather what happens when predictive inhibition is absent (Bastos et al., 2020a). So, the PR theory does not assume a distinct population of “prediction error cells” with a unique function like in PC. Instead, PR suggests certain cells become rhythmically engaged in the formation and routing of predictions.

Here, we compared the three models above against empirical LFP data. Our approach has three steps, illustrated by arrows in Figure 2. The mathematical details of our approach are explained in the *Methods* section and the Supplementary Material. Our approach includes a combination of learning (similar to neural network training) and model inference of the sort used in e.g. Dynamic Causal Modeling (DCM) and similar approaches (Pinotsis et al., 2016c, 2014). A similar approach was used in (Pinotsis and Miller, 2023) to test for ephaptic effects. Below we summarize the steps of our approach.

**Figure 2.**
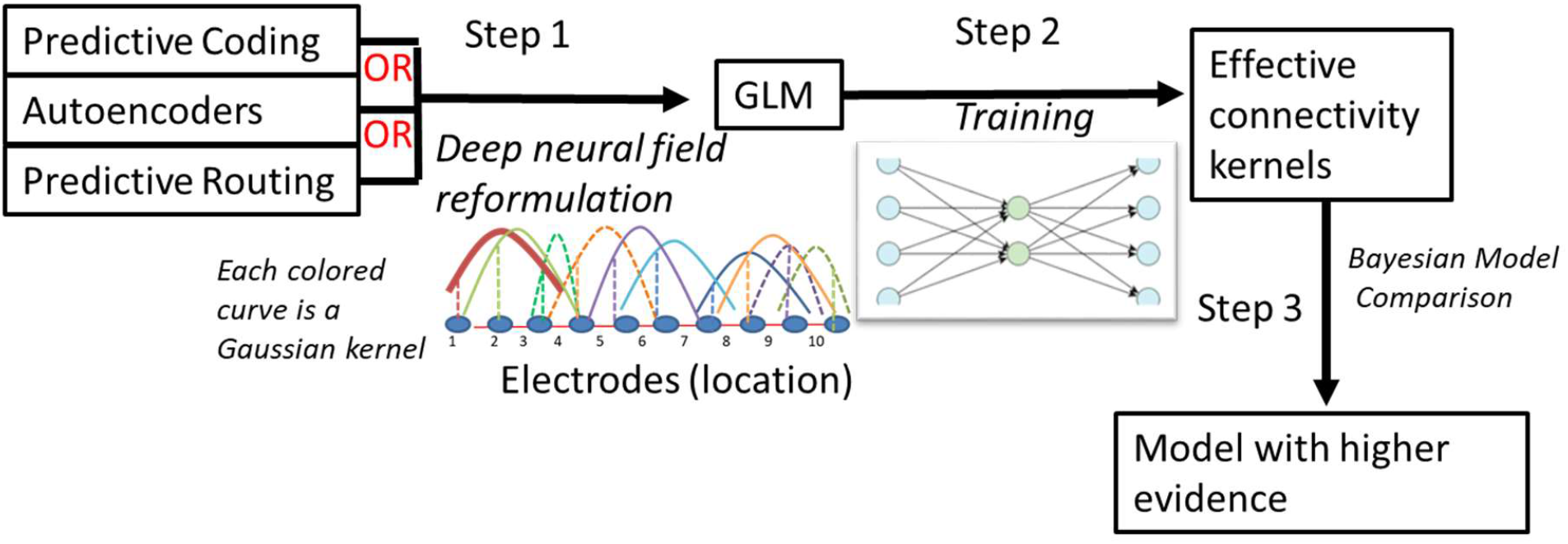
The Steps of our Approach: 1) Reformulate Predictive Coding, autoencoder networks and predictive routing as a Gaussian Linear Model (GLM) to obtain their “deep neural field” variants, see (Pinotsis et al., 2017a). 2) Train the three GLMs using LFP data from areas V4, 7A and PFC to extract the effective connectivity describing information flow in neural ensembles that might implement each algorithm. 3) Based on effective connectivity for each model, obtain predictions of neural activity and compare them against LFP data to find the most likely brain algorithm.

First, we reformulated the Predictive Coding, routing and autoencoder models as different variants of Gaussian Linear Models (GLMs; Step 1, see *Methods*). This is similar to deep neural fields used in (Pinotsis et al., 2017a). In Step 2, we performed latent state extraction from each GLM after training it using local field potentials (LFPs) recorded during a visual search task (Bastos et al., 2020a). We thus obtained the effective connectivity matrix – after assuming a Gaussian distribution (colored dash lines in Figure 2; (Pinotsis and Miller, 2022). This is similar to a connectivity matrix in classical neural networks. In Step 3, we used Bayesian Model comparison (BMC; (Friston, 2008; Kass and Raftery, 1995; Pinotsis et al., 2012), and obtained estimates of model evidence i.e. how likely is for each of the three (algorithm—specific) models –implementing Predictive Coding, autoencoder, predictive routing—to have generated the observed LFPs. We start with Steps 1 and 2 below.

We first studied differences in the organization of neural activity induced by Predictive Coding, autoencoders and predictive routing. These were quantified in terms of connectivity in cortical circuits that might be implementing each of the three brain algorithms. Since different algorithms induce a different pattern of information flow in the cortical circuit, it is reasonable to expect that the connectivity will also be different. Note that any such differences cannot be attributed to differences in the training process we used, since this is the same across all three algorithms tested: we reformulated Predictive Coding, autoencoders and predictive routing as a Gaussian Linear Model (GLM) trained with a Restricted Maximum Likelihood algorithm (*Methods*). This reformulation yields deep neural field variants of the three algorithms that contain learned weights, see (Pinotsis et al., 2017a). The output of this training included latent states that characterize the effective connectivity in the neural circuit from which the laminar electrodes record.

Here, we used LFP data recorded using laminar electrodes from supra and infragranular cortical layers in areas V4, 7A and PFC of the monkey cortex. This was the only triplet in the dataset (Bastos et al., 2020a) with laminarly resolved data. Having such data is important for testing the algorithms considered here, because they make specific assumptions about the role of deep and superficial pyramidal cell populations. LFPs are thought to describe neural activity from a population in the proximity of each electrode. Using the FLIP method (Mendoza-Halliday et al., 2024), we identified the particular electrodes where alpha and gamma activity peaks as well as the crossover electrode, where the mean relative power of alpha and gamma frequency bands intersect. Here we consider area 7A that is in the middle of the brain hierarchy. It is where the integration of top down and bottom up signals suggested by Predictive Coding would happen. LFP data from this area will be used for comparing the evidence of the three algorithms below. The results of our FLIP analyses for are 7A are shown in Figure 3A. We found that alpha activity peaked at electrode 8 and plateaued for deeper contacts (red line inside orange ellipse). The cross-over electrode was identified as electrode 7. We used LFP data from the 6 electrodes in infragranular layers (electrodes 1—6) to capture activity in error populations and another 6 below the cross-over electrode (electrodes 8—14) to capture activity in state populations, including the electrode where beta activity peaked. Brain theories like autoencoders and Predictive Coding suggest that these layers contain state representations. Electrodes above the crossover captured activity for the error population in the Predictive Coding and routing models –assuming that error representations are maintained in supra-granular layers. Electrodes were numbered in a monotonic fashion; neighbouring electrodes had adjacent numbers. Thus, as the electrode number increased, cortical depth from which the LFP data were recorded increased too.

**Figure 3.**
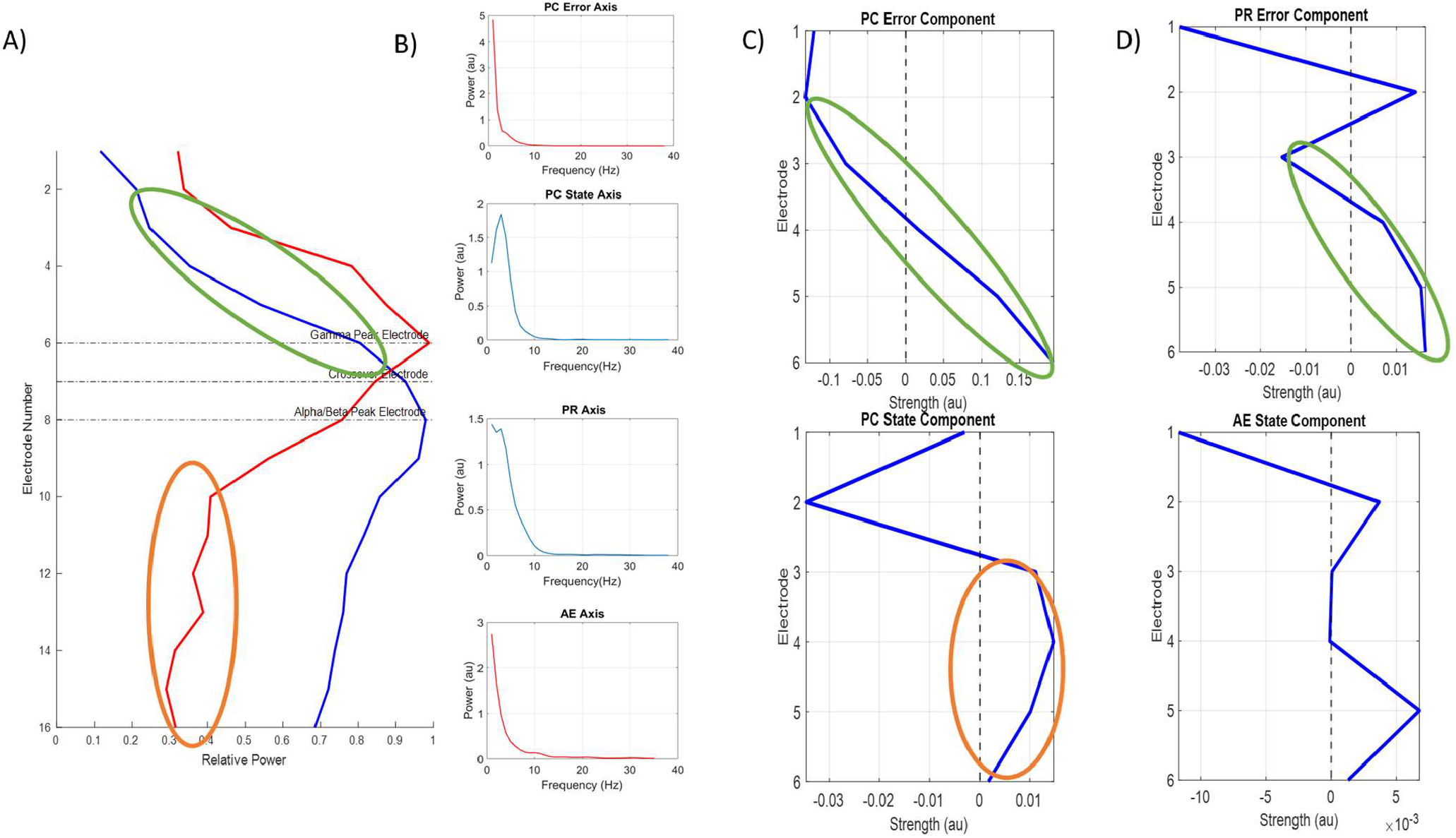
(A) Relative alpha (10—19Hz) and gamma (75—150Hz) power profiles across cortical layers. These were obtained using the FLIP method. Electrodes are shown on the vertical axis and power on the horizontal. The electrodes with maximum alpha and gamma power are shown with dashed lines, alongside the cross-over electrode where their relative power values in the two bands are equal. (B) Power spectra obtained from principal axes for error and state populations for different models. Principal axes strengths are the instantaneous scale factor with which the corresponding component strength (shown in C and D) must be multiplied to reconstruct the observed LFP. These reveal the dominant frequencies in LFP activity for each population at the largest spatial scale. (C) Connectivity components for error (top) and state (bottom) populations in the Predictive Coding model. Electrodes are shown on the vertical axis and strength on the horizontal. These are low-dimensional latent modes that capture a characteristic pattern in the effective connectivity among neural populations sampled with laminar electrodes. They help reveal the dominant connectivity motifs underlying LFPs, and link changes in network function with cognitive function. The dashed line corresponds to zero component strength (shown in x axis). Notice the similarity with the FLIP result, that is a monotonic increase in superficial layer (inside the green ellipse, see also A) and flat profile in deeper layers (inside the orange ellipse, see also A) (D) Connectivity components for predictive routing (top) and autoencoder (bottom) populations. As in C), notice the similarity of the PR component with the FLIP result in deeper layers (monotonic profile, inside the green ellipse, see also A).

Using the LFP data recorded from V4, 7A and PFC (areas *j, j+1* and *j+2* in *Methods*), and the PC model shown in Figure 1B, we obtained the connectivity components 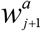 of the state population in 7A (Figure 3C). These correspond to intrinsic (within area) weights – i.e. synaptic weights in neural ensembles in an animal that has learned to perform the task. These are then used as priors for fitting the models in the model comparison below, see also (Pinotsis and Miller, 2023). To do this, we implemented Step 1 and Step 2 in Figure 2 discussed above. We constructed a GLM model from the Predictive Coding model (Equation (15) in *Methods*). We then trained this model using LFP data from deep layers in 7A and extracted its latent states. We also trained a GLM for the PC error population (Equation 16) using data from 7A superficial layers. Besides PC, we similarly trained GLMs for the predictive routing (Equation 11) and autoencoder (Equation 9) models. Details and the rationale of our approach was explained in the section *Learned connectivity in Predictive Coding* above. In brief, using a GLM model provides connectivity components that describe the spatial profile of activity predicted by the corresponding model and principal axes that describe the corresponding neural dynamics. These are shown in Figures 3B—D. Principal axes are matrices with dimensions corresponding to the number of time samples by the number of trials, representing the instantaneous contribution to the recorded LFP data averaged across electrodes. Similar to PCA axes, the first GLM axis shown here captures fluctuations in neural activity at the broadest spatial scale, allowing the identification of sparse, large-scale differences in the neural activity predicted by the three models. The first axes for the PC, PR and AE models are shown in Figure 3B. The PC axis for the error population is shown at the top followed by the PC axis for the state population, then the PR axis for the error population and the AE axis for the state population at the bottom panel – assuming that the AE model provides alternative predictions for deep layer, state activity and the PR model for superficial layer, error activity. Note that the PC error and AE axes have the usual 1/f behavior while the PC state and the PR axes peak in the theta range. All axes show significant theta and alpha activity; suggesting that these are the dominant frequencies at large spatial scales captured by the connectivity components. The strengths of connectivity components across electrodes shown on the horizontal axes of Figures 3C and D correspond to the level of activation in these frequency ranges. Large positive component strengths (on the right of the vertical dashed line in Figures 3C and 3D) suggest large LFP responses for that electrode. Recall that these are the weights with which signal is attenuated or amplified as it reaches the recording electrode from other electrodes (Pinotsis and Miller, 2023). Negative strengths suggest inhibitory drive and reduced LFP activity.

In Figures 3C and 3D, strengths are positive and negative (on the left and right of the dashed line) for different electrodes. These switches simply suggest the well known dipole structure of electrical current flow in pyramidal cells (Hämäläinen, 1987) picked up by laminar electrodes. Also, note that the PC error component shown in the top panel of Figure 3C has some similarities to the alpha power profile obtained with the FLIP method shown in Figure 3A: for electrodes 2—6 it shows a similar monotonic increase of low frequency activity in superficial layers (inside the green ellipses in Figures 3A and 3C). The PR axis shown in the top panel of Figure 3D also shows a similar monotonic profile (green ellipse). The PC state component shown in the bottom panel of Figure 3C shows a similar flat profile of low frequency activity to what was obtained with FLIP for electrodes 10—13 in Figure 3A, see orange ellipses, while the AE axis in the top panel did not show the flat profile observed in the PC state component and FLIP recordings.

To confirm the connectivity components found above, we followed the approach by (Pinotsis and Miller, 2022) and compared them with clusters found using the method of (Humphries, 2011). Recall that the connectivity components describe scaling factors (weights) by which signal incoming to a population near an electrode is weighted by (Pinotsis and Miller, 2023). The method of (Humphries, 2011) provides an alternative way to identify connectivity patterns through unsupervised clustering. We adapted the original algorithm to process (LFP) data instead of spike trains. For each trial, the method assigned an ensemble index to each electrode—effectively clustering electrodes into ensembles (groups of electrodes with similar activity). To sum, the method by (Humphries, 2011) finds effective connectivity in terms of cluster indices. To test whether this confirms our results above, we computed the correlation between connectivity components and cluster indices. These are shown in Supplementary Figure 1C. Figures S1A and S1C show the correlation coefficients for PC and PR models, while Figures S1B and S1D the corresponding *p*-values. Connectivity strengths show high correlations (*R*^*2*^>0.8) and are significant for most of the trials (shown with green bars).

The connectivity components characterize the overall structure of brain responses –found above to reflect a dipole source, as expected. However, the connectivity matrices, not the components used above, describe the usual pairwise interactions between electrodes. These describe how signal propagating across different 7A cortical depths, sampled with laminar electrodes is amplified or attenuated. Connectivity matrices can be readily obtained from the components assuming a Gaussian connectivity profile over space (coloured lines in Figure 2; Pinotsis and Miller, 2022). A Gaussian profile is a common assumption that describes a connection probability that drops off with distance (Bullmore and Sporns, 2012; Coombes, 2005). The connectivity matrices are of dimension *N*_*s*_-by-*N*_*s*_ (square matrices), where *N*_*s*_ is equal to the number of electrodes. They are shown in Figure 4.

**Figure 4.**
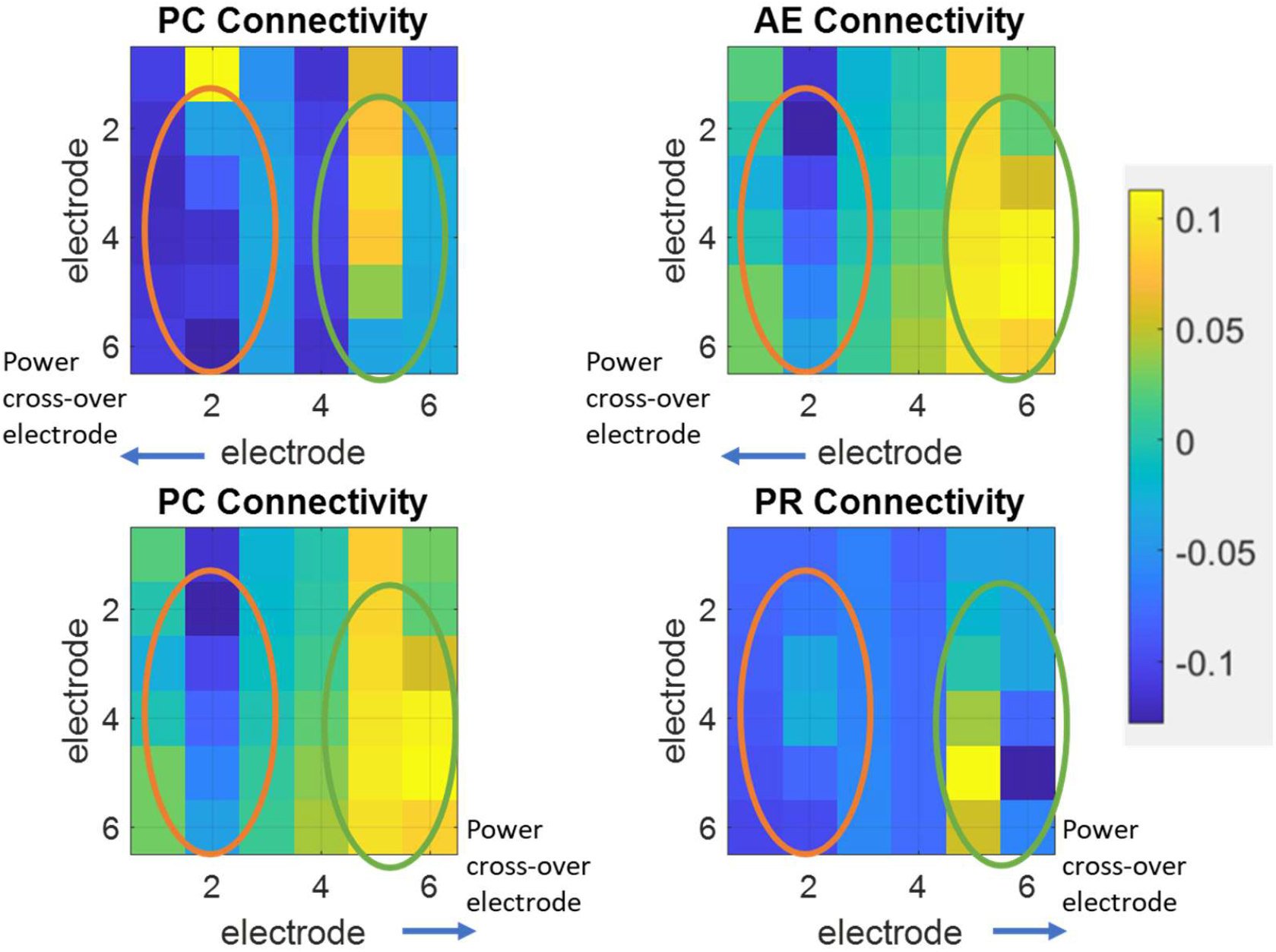
Top row: Connectivity kernels for deep layers in the Predictive Coding model (A) and the autoencoder model (B). Each entry corresponds to the weight that electrical signal is multiplied with (i.e. it is amplified or attenuated) while propagating from the row to the column electrode. Overall, results were similar across models. In deep layers, both the PC and AE kernels predict inhibitory input near the power cross-over electrode (blue entries dominate inside the orange ellipses in panels A and B). Polarity is reversed at the other end of the laminar electrode array (green ellipses in the same panels). Bottom row: Connectivity kernels for superficial layers in the Predictive Coding model (C) and the predictive routing model (D). Again, both models make similar predictions. In superficial layers, both the PC and PR kernels predict excitatory drives near the power cross over electrode (green ellipses in panels C and D). Polarity is again reversed at the other end (orange ellipses).

In the matrices shown in Figure 4, each entry corresponds to the weight of interaction between populations and the electrodes that sample from them (e.g. entry (1,2) determines the interaction between electrodes 1 and 2). In other words, signal leaving electrode 2 reaches electrode 1 after being weighted by the value in entry (1,2), similar to connection weights in a neural network. For state populations in deep layers, both models (Figure 4A and 4B) predict inhibitory input near the cross-over electrode (orange ellipses). They predict inhibitory recurrent connectivity from layer 6 to layer 4 electrodes that has also been linked to eliciting action potentials in output layer 5 (Kim et al., 2014). Also, both models predict deep layer excitation of output layers (green ellipses in Figures 4A and 4B; Kim et al., 2014). For populations in superficial layers, both the PC and PR models make similar predictions (Figures 4C and 4D): local recurrent excitation around input layer 3 possibly as a result of feedforward drive (green ellipses; Ungerleider et al., 2008). Also, both models predict weak recurrent inhibition in superficial layers (orange ellipses in Figures 4C and 4D; Scholl et al., 2019) that has been observed in layers 2/3 of the visual hierarchy as a result of normalization and gain control (Bastos et al., 2012).

### Predictive Coding vs. alternative models

The effective connectivity matrices describe the information flow between laminar electrodes during the visual search task considered here (Bastos et al., 2020a). Knowing these matrices, we can obtain predictions of neural activity in space and time –similarly to knowing the connectivity in a neural network (*Methods)*, see also Supplementary Figure 2A. Based on model predictions and LFP data, we can compare alternative models (Step 3 in Figure 2). We computed the Free energy for each model and trial. We compared the AE and PC models using LFP data from the contacts sampling deep layers and the PR and PC models using superficial LFP data. We used a Bayesian algorithm that allowed us to obtain an approximation of model evidence, that is, how likely the data is given a specific model that might have generated them. This approximation to model evidence is called Free Energy. This is a useful measure from Bayesian inference that allows us to assess which of alternative models might explain the data better. We used Free Energy and Bayesian Model Comparison (BMC; see Pinotsis et al., 2018 for details) to assess the relative evidence of various models. The advantage of the Free Energy approximation is that it balances model fit and complexity: under a maximum likelihood estimation (which provides the best fitting parameters), this cost function penalizes overly complex models. Such models include parameters that do not explain away distinct parts of the data. Thus, Free Energy favours simpler models (cf. Ockham’s razor). In brief, BMC involves fitting competing models to LFP data to determine the most likely model. The output of BMC is an odds ratio known as the Bayes Factor (BF; Kass and Raftery, 1995). The Bayes Factor is a Bayesian analogue of the traditional odds ratio, quantifying the probability that a particular model, among several alternatives, best explains the LFP data. Here, it provides an estimate of how more likely a given model (algorithm) might be to support brain computations during an attention task compared to other models. In the case of two competing models, this is simply the difference of the Free Energy approximation to model evidence.

In our first model comparison, we considered LFP data from the superficial layers of area 7A. These receive feedback input from PFC. We compared the PC and PR models. This analysis assesses whether errors,that is differences between feedback signals carrying predictions and feedforward signals carrying sensory input are explicitly computed by the brain, like the PC model proposes, or not –like the PR model suggests. We analysed three sessions using different data from the data used for training the models above to avoid data leakage. Example results of this analysis are shown in Supplementary Figure 2B. The panel shows a comparison between the PC model for error representations against the PR model obtained by fitting both models to LFP data from superficial cortical layers. Here, trials from a single session are shown. The fixed effects Bayes factor was *BF*≈*982* in favour of the PR model. This suggests that predictive suppression from deep layers is sufficient to explain superficial layer activity, without the need to explicitly compute error signals. We repeated the analysis for two other sessions and again found that the PR model was more likely in area 7A with Bayes factors *BF*≈*1092* and *BF*≈*1109*. This lends support to predictive routing as refined version of Predictive Coding to which we will come back in the Discussion.

We then considered state representations and data from deep cortical layers in area 7A. These receive feedforward input from V4. Model comparison involved the PC and AE models. The alternative hypotheses included whether state representations follow solely from feedforward signals—like the AE model proposes, or the combination of feedforward and feedback signals –as the PC model suggests. Here, Bayes factors suggested that the PC model was more likely than the AE model for all three sessions and three areas considered. By averaging differences after subtracting the Free Energy estimates, we obtained the mixed effect Bayes factors as before. We found the following Bayes factors for the three sessions, *BF*≈*175, BF*≈*164*, and *BF*≈*152* in favour of the PC model. The results of our analysis suggest that Predictive Coding is more likely than autoencoders to be implemented by the brain.

## Discussion

We compared different algorithms the brain might be using for computations: Predictive Coding, routing and autoencoders. We reformulated neural network implementations of these algorithms as Gaussian Linear Models (GLMs) and then fit these models to LFP data from a visual search task (Bastos et al., 2020a). We found that a combination of Predictive Coding and routing models had the highest evidence with this data: State representations in deep layers seemed to require hierarchical message passing from areas above and below like Predictive Coding suggests; at the same time, no explicit error computations seemed to take place in superficial layers and predictive suppression from deep layers offered a simpler explanation –as it follows from predictive routing.

Predictive Coding (PC) proposes that the brain operates within a hierarchical framework (Friston, 2010; Hohwy et al., 2008; Kanai et al., 2015). This view is consistent with anatomical and functional studies (Bastos et al., 2012; Douglas and Martin, 2004; Haeusler and Maass, 2007), and implies that the brain’s organization not only reflects structural connectivity but also constrains how learning occurs. Earlier work likewise emphasized the hierarchical arrangement of cortical networks (Felleman and Van Essen, 1991; Lynn and Bassett, 2019; Markov and Kennedy, 2013). In PC, sensory input is passed in a feedforward manner from lower to higher cortical areas, with each level transforming and relaying information from the one below. At the same time, reciprocal connectivity ensures the presence of feedback signals, flowing from higher to lower areas. Feedforward information is conveyed by supragranular neurons, while feedback is carried by infragranular neurons (Lamme et al., 1998; Markov et al., 2013).

In Predictive Coding, feedforward signalling relies on AMPA receptors (Rivadulla et al., 2001) transmitting prediction errors, but these signals are strongly regulated by local inhibitory interneurons (Shipp et al., 2013). We found such excitatory layer 3 inputs and inhibitory regulation here, similarly to (Ungerleider et al., 2008). Our PC model identified inhibitory recurrent connectivity from layer 6 to layer 4 (Kim et al., 2014) and local inhibitory inputs in superficial layers possibly suppressing redundant or fully explained activity (Rao and Ballard, 1999). Another feature of the PC model is that inhibition acts as a dynamic filter that shapes error signaling and maintains the stability of hierarchical inference. In our formulation, Predictive Coding uses dynamic constraints to solve the inverse problem of reconstructing recognition density (Friston, 2010; Pinotsis et al., 2016b). Thus, optimization was carried out in an input-specific manner rather than through a generic representational scheme considered in alternative theories. The structural and functional organization with feedforward and feedback connections also provides a substrate for PC’s central claim: that perception and cognition emerge from learning at each hierarchical level. At the same time, feedback and errors are not unique to PC, as it is also integral to other theoretical accounts such as Temporal Difference learning (O’Doherty et al., 2003) or Reinforcement Learning (Sutton and Barto, 1999). However, PC assigns a distinct role to feedback: it carries predictions, shaped by prior experience, that act as hypotheses about incoming feedforward signals. In this view, conscious perception arises when predictions are confirmed by sensory input, aligning expectations with evidence.

Predictive Routing (PR) is a recent alternative to PC (Bastos et al., 2020a). Here we have assumed that PR compresses sensations into independent hidden variables, uses them to predict the data, and learns representations that keep as much useful information as possible—while Predictive Coding instead relies on learned expectations from experience. Mathematically, we implemented a bottleneck architecture with independent Gaussian priors, unlike Predictive Coding that employs empirical priors. In PR, recognition is implemented by connections from latent states to predictions similarly to an information bottleneck (Tishby et al., 2000). This mirrors an important difference in how inference is performed by the underlying brain circuitry: PR argues that feedback primarily regulates the excitability and routing of signals, without requiring explicit representational content from other areas. In PC, conscious perception arises when feedforward signals confirm feedback “guesses,” whereas in PR, perception reflects the selective amplification (or inhibition) of relevant (or irrelevant) feedforward pathways.

Anatomically, both PC and PR draw on the distinct laminar and frequency-specific organization of feedforward versus feedback connections (Shipp, 2016), but PC emphasizes representational precision-weighting and Bayesian inference, while PR emphasizes dynamic gain control and oscillatory routing. Thus, layer 2/3 activity is not a specialized error signal, but rather what happens when predictive inhibition is absent (Bastos et al., 2020a). In this view, the routing of information is biased to enhance relevant inputs while suppressing irrelevant ones (Gabhart et al., 2023). PR shares a bottleneck architecture with a popular algorithm known as autoencoder (AE; Hinton and Salakhutdinov, 2006). This is another alternative to PC. The bottleneck architecture forces both PR and AE algorithms to capture only the most informative statistical structure of the LFPs.

Implementing the PR model here, we obtained similar predictions to the PC model with regard to recurrent inhibition effects in superficial layers and excitatory drive in input layers. Similarly, implementing the AE model, we obtained similar predictions to the PC model with regard to deep layer 6 inhibitory drive to layer 4 neurons (Kim et al., 2014) Thus, to assess which of the models was more likely, we compared the evidence of these models against LFP data.

We used a combination of learning and inference similar to (Pinotsis and Miller, 2023). The models first learned the connectivity parameters in neural ensembles in each area (V7, 7A and PFC) of the brain hierarchy –corresponding to synaptic weights in an animal that has learned to perform the task. These were subsequently used as priors in a model comparison that assessed the evidence against LFP data by focusing on the middle area 7A. This is where the integration of top down and bottom up signals suggested by Predictive Coding would happen. We compared PC vs. AE in deep layers and PC vs. PR in superficial layers. Our assumption was that Autoencoders predict state activity corresponding to beta oscillations in deep layers (Pinotsis et al., 2018; Pinotsis and Miller, 2020) while Predictive Routing predicts superficial gamma activity without requiring an explicit error representation (Bastos et al., 2020a). Using Bayesian Model Comparison (Kass and Raftery, 1995), we compared the relative evidence of these models. After calculating a Free energy measure of approximate evidence, we found that the PC algorithm was more likely than the AE algorithm in deep 7A layers but, interestingly, less likely than the PR algorithm in superficial 7A layers. This points to an emergence of state representations from hierarchical message passing between brain areas in prediction-based circuits that constantly refine their internal models of the world in deep layers, but do not require an explicit recursive error correction in superficial layers. Thus, Predictive Coding seemed to explain well the formation of state representations in deep cortical layers as a result of hierarchical message passing, while predictive routing offers a simpler explanation of feedforward signals in superficial layers in terms of local predictive suppression from deep layers –unaffected from feedback input from the cortical hierarchy.

Since the introduction of Marr’s three levels of analysis (Marr, 2010)—computation, algorithm, and implementation—in the study of information-processing systems, significant progress has been made in identifying the neural circuits responsible for implementing brain computations (Maass et al., 2004; Niell and Scanziani, 2021). Dissecting the anatomical and functional organization of these circuits has been possible following advances in neuroimaging techniques, such as electrophysiology, optogenetics, and calcium imaging (Deisseroth, 2011; Ghosh et al., 2022; Michel and Murray, 2012; Susaki et al., 2014). At the same time, new theories have been put forward that allow one to hypothesize which algorithms the brain might be using like the ones considered here (Baldi, 2012; Bastos et al., 2012; Bastos et al., 2020b; Girin et al., 2020; Pinotsis et al., 2018; Rao and Ballard, 1999; Whittington and Bogacz, 2017).

Recent advances in brain imaging allow one to test brain theories. For instance, studies using two-photon imaging have revealed the intricate connectivity patterns of cortical microcircuits that underlie orientation selectivity in the visual cortex (Barson et al., 2020). These provide experimental support for the emergence of neuronal receptive fields that was predicted long ago by efficient coding (Olshausen and Field, 1996). Similarly, advancements in connectomics have mapped the wiring diagrams of entire neural systems, such as the connectome of *C. elegans* or detailed cortical networks in mammals (Shapson-Coe et al., 2024), where connectivity can be predicted by Predictive Coding (Wright and Bourke, 2021).

Still, determining the algorithm the brain might be using to perform computations is an open challenge. To address this challenge, biologically inspired computational models try to simulate brain activity by incorporating structural and functional constraints that follow from real circuits. They try to replicate measurable behavior and observed responses and are often evaluated against empirical data using various approaches, like Dynamic Causal Modeling (Bernal-Casas et al., 2017; Pinotsis et al., 2012), Bayesian inference (Knill and Pouget, 2004), compartmental and spiking networks (Doruk and Abosharb, 2019; Reneaux et al., 2025; Rodriguez and Tuckwell, 1996) or deep neural networks (Kietzmann et al., 2017) and hybrid models (i.e. combinations of the above; e.g. Pathak et al., 2022). Such models are approximations of real circuits, trying to describe what is computed (“goal”) or the algorithm the brain might be implementing (“how”), or a combination of the two, like Predictive Coding (Hohwy et al., 2008; Rao and Ballard, 1999) and Reinforcement Learning models (O’Doherty et al., 2003; Seymour et al., 2004) and then assessing their evidence in the light of empirical data.

Even the most accurate biologically inspired models have limitations. An important one is that different brain algorithms can sometimes account for similar behaviors. Thus, accurate model predictions do not guarantee that the algorithm implemented by the model is also used by the brain. For example, model-free RL algorithms rely on cached values of past experience, whereas model-based algorithms simulate future outcomes using internal models (Dayan and Daw, 2008). Both sorts of models can accurately explain behavior, however they make divergent predictions regarding flexible generalization and adapting to new contingencies. Thus, to pin down the exact algorithm one needs to connect algorithmic theories with biophysical substrates and neural dynamics (Daw et al., 2011; Wunderlich et al., 2012). This is what we presented here.

To sum, we tested different algorithms the brain might be using to compute behavioral outputs during a visual search task. This required linking brain theories with circuit motifs in cortical areas. Different motifs led to different model predictions that were fit to LFP data to find the most likely model. This turned out to be a combination of Predictive Coding and routing models. We hope our work may contribute to bridging algorithmic theories with biophysical substrates and more generally, in understanding how the brain implements computations and in building better AI systems that rely on attention like transformers (Khan et al., 2022) and recommenders (Zhou et al., 2018).

## Acknowledgements

This work is supported by the UK Medical Research Council (MRC) (Grant Number: MR/W011751/1), Office of Naval Research (Grant Number: MURI N00014-23-1-2768), The Simons Center for the Social Brain, Freedom Together Foundation and The Picower Institute for Learning and Memory.

## Data Availability

No new data were generated during this study.

## Supplementary Figures

**Figure S1.**
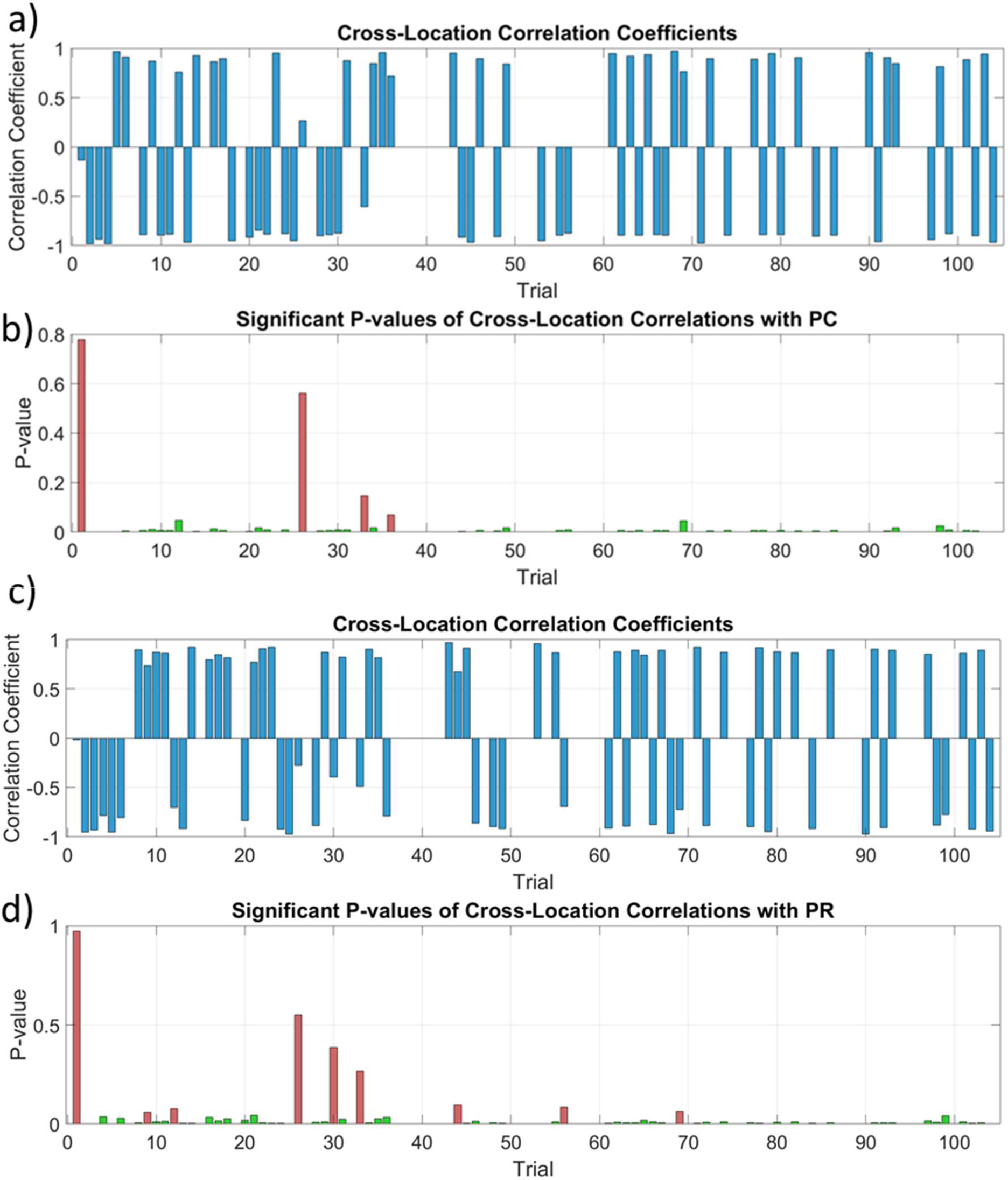
Correlation strengths (a) and corresponding p-values (b) for correlations between connectivity components of the PC model cluster indices obtained using the method of (Humphries, 2011). We found high correlations (R^2^>0.8) and significant p-values (p=10^−4^ – 10^−2^) for most trials considered. Significant trials are shown with green bars. Panels (c) and (c) show the corresponding results for the PR model.

**Figure S2.**
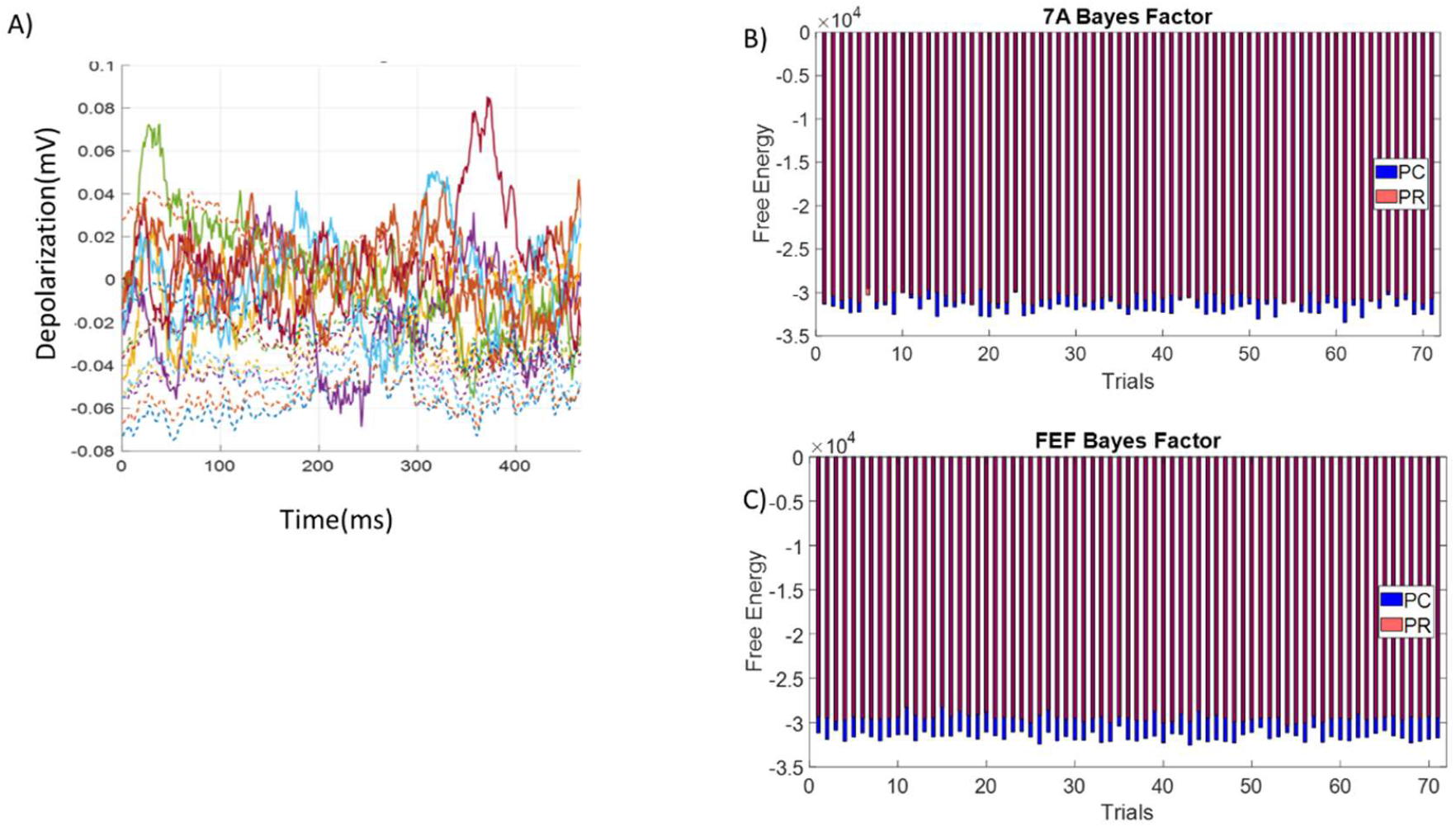
(A) Predictive Coding model predictions (solid lines) and LFP data (dash lines) across electrodes (locations) in layers 4—6. Each color depicts a different electrode. These are time series for 460ms of the attention epoch. (B) Bayes factor comparison of the PC and PR models across trials. The plot shows Free energy estimates for model comparison for two competing models, Predictive Coding and routing using LFP data from superficial cortical layers. Larger (less negative) values indicate better model evidence. Each point represents the free energy of a model on a single trial, allowing for direct comparison of model performance over time. This analysis illustrates the relative fit of the models across the experimental timeline. Note the blue lines (PC model approximate evidence) are longer than the red (PR model approximate evidence). Bayes factors (BFs) computed for each trial, quantifying the evidence in favour of the PR model over the PC model for superficial layer activity in area 7A. A fixed effects analysis yields a Bayes factor equal to 982. (C) Similar to (B) for superficial layer activity from area FE. The fixed effects Bayes Factor is equal to 2203 in favour of the PR model.

## Notes

### Competing Interest Statement

The authors have declared no competing interest.

### Summary of Updates

small typos in intro and abstract corrected

